# Artificial light at night intensifies effects of a parasitic flatworm on the water flea *Daphnia magna*

**DOI:** 10.1101/2025.02.21.639422

**Authors:** Nedim Tüzün, Franz Hölker, Luc De Meester

## Abstract

Artificial light at night (ALAN) can strongly alter organismal traits, but its role in shaping species interactions remains poorly understood, especially so in aquatic ecosystems. By capitalizing on a recently discovered antagonistic interaction between a brood-parasitic flatworm and *Daphnia magna* water fleas, we tested whether this interaction depends on exposure to ALAN. During a 19-day laboratory population growth experiment, we manipulated flatworm presence and nighttime light conditions in a full-factorial design. We confirmed the negative effects of flatworm predation on *Daphnia* abundance at the population level. Importantly, we showed that the flatworm-caused decrease in *Daphnia* abundance under ALAN (81%) was twice as strong compared to dark-night conditions (39%). Our findings are relevant for assessing the impact of ALAN on the development of *Daphnia* populations and thus top-down control of phytoplankton. Freshwater ecosystems in urbanized areas, where this parasitic interaction was first encountered, may be especially at risk, as these are typically exposed to high levels of stress factors, including light pollution.

## INTRODUCTION

Human induced environmental change shapes not only organismal traits, but also species interactions (Guiden et al. 2019). Artificial light at night (ALAN) is a pervasive stressor for a wide range of taxa, influencing life-history, behaviour and physiology, which in turn influence species interactions and ultimately community structure and ecosystem functions (Gaston et al. 2015; Sanders et al. 2021, Hölker et al. 2021). Whether and how species interactions are shaped by ALAN has recently gained attention among ecologists (Seymoure et al. 2023). Aquatic habitats are suggested to be particularly threatened by ALAN, because aquatic organisms often have reduced available refuge to avoid light exposure, especially so in urban ponds with limited structural complexity (Oertli & Parris 2019) and are expected to be exposed to high levels of ALAN (Hölker et al. 2023). Despite this, effects of ALAN on species interactions in aquatic ecosystems remain understudied (Hölker et al. 2023).

Natural temporal light pattern produced by nocturnal celestial bodies are important environmental cues for many animals, including aquatic species (Kühne et al. 2021, Marangoni et al. 2022, Ganguly & Candolin 2023). For example, moonlight can induce the vertical migration of zooplankton down to a depth of 100 m (Last et al. 2016). The high sensitivity of aquatic organisms to low-intensity natural light makes them susceptible to disturbance even by low-intensity ALAN (Hölker et al. 2023). It is known that ALAN interrupts natural diel vertical migration patterns (Moore et al. 2000, Ludvigsen et al. 2018, Maszczyk et al. 2021), where zooplankton species stay in deep waters during the day to minimize the risk of predation by visual planktivorous fish and only ascend to the surface water layers at night to feed on phytoplankton and microzooplankton (De Meester et al. 2022). Disruptions of diel activity and habitat selection patterns can result in altered temporal niche partitioning, which in turn can influence species interactions, for instance resulting in increased predation rates when interactors occupy the same temporal niche (Seymoure et al. 2023). Aside from changed activity patterns, ALAN may also alter species interactions by imposing physiological stress on the interacting organisms (e.g., Grubisic et al. 2019; Ganguly & Candolin 2023).

To test whether ALAN has an influence on species interactions in aquatic ecosystems, we capitalized on a recently discovered interaction between a flatworm and *Daphnia* water fleas (Tüzün et al. 2025). The water flea *Daphnia*, a key ecological interactor in freshwater ecosystems, is well suited for this question. First, they show a changed pattern of diel vertical migration under ALAN, with the animals staying close to the bottom during both day and night (Moore et al. 2000, Maszczyk et al. 2021). Second, *Daphnia* have numerous antagonistic interactions with a wide range of organisms. We have recently shown that the typhloplanid flatworm *Strongylostoma simplex*, an egg parasite of *Daphnia* (Fig. 1), has negative effects on short-term individual *Daphnia* performance in terms of survival and offspring production (Tüzün et al. 2025). Here, we first aim to test the hypothesis that the effects of the flatworm on individual *Daphnia* translates into changes at the population level. We also aim to test the hypothesis that ALAN significantly influences the effects of flatworms on *Daphnia* abundance. While little is known about the diel vertical migration patterns of flatworms (e.g. De Meester & Dumont 1990), we predict the low depth preference of *Daphnia* under ALAN to result in an increased encounter rate between *Daphnia* and flatworms (assuming the flatworm is benthic: Dumont et al. 2014). We therefore expected a reduced *Daphnia* abundance under ALAN in the presence of flatworms.

**Figure 1.**
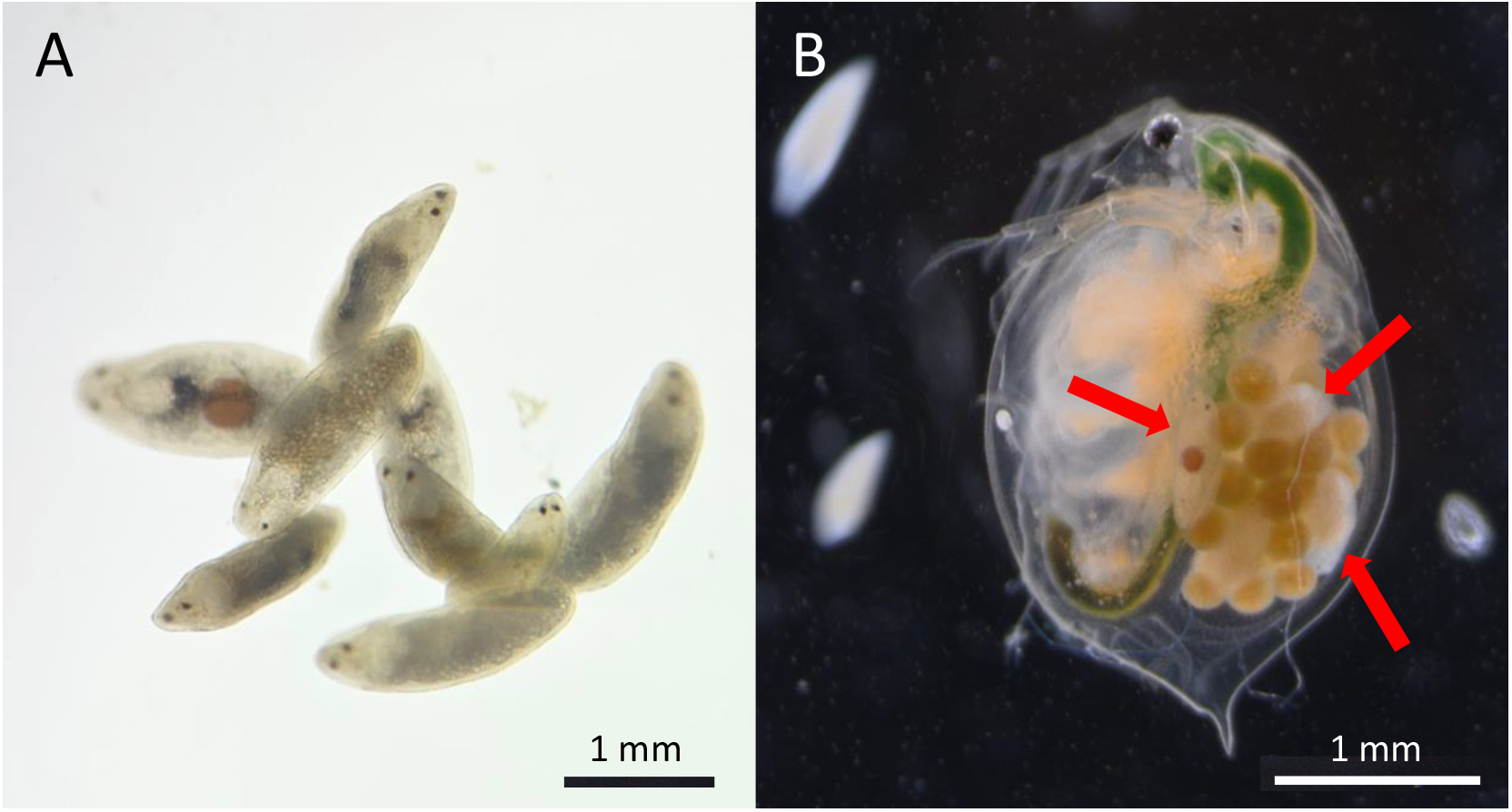
The typhloplanid flatworm *Strongylostoma simplex* (A) and the water flea *Daphnia magna* (B). In panel B, flatworms inside the brood chamber of a water flea with eggs are indicated with arrows. Note also on the left side of the water flea the two free swimming flatworms. Photo credit for panel A: Mareike Brehm-Benedix.

## MATERIALS AND METHODS

### Experimental settings

To explore the effects of flatworms on *Daphnia* abundance under artificial light at night (ALAN), we conducted a population growth experiment over 19 days. The experimental design consisted of a flatworm treatment (flatworm absent / present) crossed with an ALAN treatment (dark at night / light at night), using four replicates per treatment combination (N = 24). Treatments were assigned to experimental units using a randomization procedure.

To produce the light environment during day and night, we used LED lamps (Sylvania L300, 3000K, warm-white LED, Feilo Sylvania International Group, Hungary), and used neutral density filter foil (Reinan, USA) wrapped around the lamps to dim the light intensity for the ALAN treatment. The illuminance during day hours was ca. 400 lx (ca. 6.4 µmol photons m^2^ s^1^ as measured on top of the jars with a sensitive illuminance meter; ILT-1600, International Light Technologies, USA). For the ALAN treatment, illuminance during night hours was ca. 35 lx (ca. 0.5 µmol photons m^2^ s^1^), compared to <0.02 lx (ca. 0.0004 µmol photons m^2^ s^1^) for the control treatment. This illuminance level is within the typically range of recorded ALAN in urban areas, albeit on the high side of that range (Hänel et al. 2018).

Jars assigned to the flatworm treatment received five individual flatworms of similar size (ca. 0.5 mm) at the beginning of the experiment. Flatworms were collected from a small artificial water body in a cemetery in Berlin, and were identified as *Strongylostoma simplex* (Tüzün et al. 2025). We used 1-l cylindrical glass jars (84 mm diameter, 210 mm height, WECK, Germany) filled with 750 ml of dechlorinated tap water. Each experimental population was started with seven *Daphnia magna* individuals of mixed age: one adult, one subadult, and five newly born juveniles. We used a single *D. magna* clone for this experiment, collected in 2022 from a very small artificial water pond in a cemetery in Berlin (52°31’19.6”N 13°30’55.6”E). The clone was kept in culture under standardized conditions (20 °C, 14:10 light:dark photoperiod) in the laboratory of IGB Berlin (clone code: ZEN-4). The habitat from which the *Daphnia* clone was isolated did not contain the flatworm at the moment of isolation. The experimental populations were fed daily with the green algae *Acutodesmus obliquus* at 1 mg C/L. We refreshed the medium (dechlorinated tap water) three times per week.

The experiment was run in a temperature-controlled incubator (Pol-Eko ST3 Smart, Poland) at 20 °C with a photoperiod of 14:10 light:dark (typical for August in Berlin). The experiment ran for 19 days, during which we counted the number of *Daphnia* three times per week. When counting the *Daphnia*, we also checked the number of worms in each jar and added new flatworms when necessary.

### Statistical analyses

To test for the effects of flatworms on *Daphnia* abundance in the absence and presence of nighttime light pollution, we constructed a linear model with time of the experiment (continuous), flatworm treatment (categorical: flatworms absent / present), and light pollution treatment (categorical: dark at night / light at night) as fixed effects. We included all interaction terms of these fixed effects in the model. Given the typical non-linear shape of *Daphnia* abundance over time, we fitted a second-order polynomial function of time. In addition, we fitted logistic growth curves to test for differences in carrying capacity and growth rate during the exponential phase (note that logistic growth models would not converge for the flatworm treatment groups due to not reaching a plateau [see Fig. 2], and were therefore excluded from this analysis). Pairwise comparisons of treatment-effects and regression slopes using estimated marginal means were calculated using the R package *emmeans* (Lenth 2023). Logistic growth models were fitted using the R package *nlme* (Pinheiro et al. 2023), and the “reduced” (no treatment differences in logistic curve parameters) vs “full” (treatment-specific logistic curve parameters) models were compared using a likelihood ratio test. All analyses were performed in R version 4.3.2 (R Core Team, 2023).

**Figure 2.**
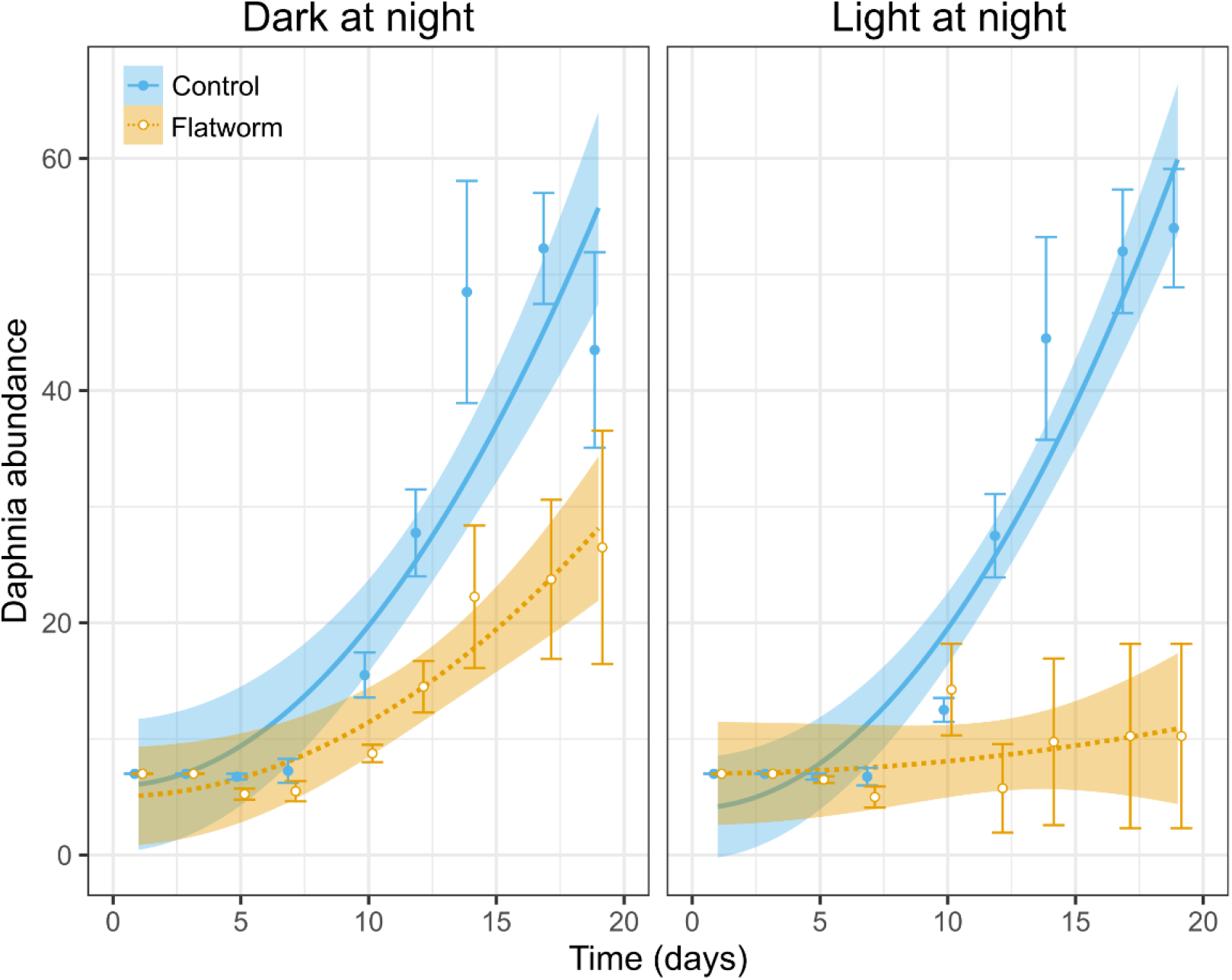
*Daphnia* abundance over time as a function of the presence or absence of predatory flatworms and the presence or absence of light pollution at night. Left panel represents no light pollution (i.e. dark at night) condition, right panel represents light pollution condition. Shown are regression lines for the control (blue, full line) and flatworm treatments (yellow, dashed line). Shown are also raw means (± 1SE) per time point. Bands around lines represent 95% confidence intervals (derived from the linear model; see Table 1). Logistic growth curves fitted for control treatments are shown in Fig. S1.

## RESULTS

*Daphnia* abundance on average increased over time, with a steeper increase after the relatively stable first week (indicated by the significant quadratic term of time, Table 1, Fig. 2). While the flatworm treatment caused on average a decrease in the *Daphnia* abundance, this decrease was more pronounced after the first week of the experiment (time^2^ × flatworm treatment, Table 1, Fig. 2). The significant three-way interaction further shows that in the light pollution treatment, the flatworm-caused decrease in *Daphnia* abundance was more pronounced compared to the control light conditions, especially toward the end of the experiment (Table 1, Fig. 2). This is further reflected as a significant difference between the linear growth rates (i.e. slopes) of flatworm-exposed *Daphnia* at control vs light pollution conditions (contrast test: t ratio = 2.59, *p* = 0.011). Considering only the no-flatworm treatments, the logistic growth curves did not differ between the control and light pollution treatments (LRT: ratio = 1.71, *p* = 0.634, Fig. S1); this is reflected in the similar values for the three logistic curve parameters across light pollution treatments (Table S1).

**Table 1.** Result of the linear model testing for effects of time, flatworm treatment, light pollution treatment, as well as their interactions, on *Daphnia* abundance. Note that time was included as a second-order polynomial term (implemented using the *poly(x,2)* function in R).

| Fixed effects | df | F value | *p* |
| --- | --- | --- | --- |
| Time<sup>2</sup> | 2 | 97.44 | <0.001 |
| Flatworm treatment | 1 | 67.83 | <0.001 |
| Light pollution treatment | 1 | 2.15 | 0.145 |
| Time<sup>2</sup> × Flatworm | 2 | 34.41 | <0.001 |
| Time<sup>2</sup> × Light pollution | 2 | 1.04 | 0.357 |
| Flatworm × Light pollution | 1 | 2.69 | 0.103 |
| Time<sup>2</sup> × Flatworm × Light pollution | 2 | 3.97 | 0.021 |

## DISCUSSION

Our results show that artificial light at night (ALAN) can exacerbate the adverse effects of brood-parasitic flatworm on *Daphnia magna* water fleas. The interaction between these *Strongylostoma* and *Daphnia* was recently described for the first time, with observations clearly suggesting an egg parasitism behaviour of the flatworm, with first indications that survival and offspring production of *D. magna* can be strongly reduced in the presence of flatworms (Tüzün et al. 2025). In the current study, we indeed confirm that this impact has negative effects on population development.

Importantly, we show that the reduction in *Daphnia* abundance caused by flatworm parasitism interacts synergistically with ALAN, i.e. it is twice as pronounced under ALAN when compared to control light conditions (final population size reduced by 81% vs 39%, respectively). Our findings have relevance for assessing the impact of ALAN on the development of *Daphnia* populations and thus top-down control of phytoplankton in standing freshwater ecosystems, especially in urban areas, as these are typically exposed to high levels of light pollution (Hölker et al. 2023).

While organismal responses to ALAN are well-documented (Hölker et al. 2021, Sanders et al. 2021), there have been much less studies on species interactions under ALAN (Seymoure et al. 2023, Hirt et al. 2023). Light pollution can shape species interactions amongst others by altering behaviour and physiology in a way that affects the encounter rate between the interactors, as illustrated for host-parasite interactions in aquatic systems (Poulin 2023). Fish predation, which also induces zooplankton to reside at greater depths in the water column, has been linked to increased parasitic infection in *Daphnia* due to increased exposure to parasite spores (Decaestecker et al. 2002). Most typhloplanid flatworms prefer benthic habitats (Dumont et al. 2014), which may explain our observation that the flatworms have a stronger impact under ALAN. An alternative pathway is that ALAN-induced physiological stress (e.g. in phytoplankton: Diamantopoulou et al. 2021; in *Daphnia*: Li et al. 2022; in fish: Kupprat et al. 2020) may have an effect on resource allocation, potentially making *Daphnia* more vulnerable to flatworm parasitism.

Increased (egg) predation pressure by flatworms under ALAN can potentially strongly impact population dynamics of *Daphnia*, given that typhoplanid flatworms can be important predators of *Daphnia* (Dumont et al. 2014). This may not only affect *Daphnia* densities and top-down control of phytoplankton, but may also lead to profound changes in zooplankton community composition (e.g. Devkota et al. 2023). This is because *S. simplex* is a parasite of eggs that are in the brood chamber of *Daphnia*, which likely makes large-bodied species more vulnerable than smaller ones (as shown for copepods predating on *Daphnia* eggs, Gliwicz & Lampert 1994). Increased fish predation on zooplankton under ALAN, for example, reduced the mean body size of zooplankton and changed the zooplankton community structure (Tałanda et al. 2022). ALAN effects on the flatworm – *Daphnia* interaction may have similar effects.

The effects of ALAN on species interactions and ecosystem functions may be further exacerbated by other stressors, a key one in an urbanization and climate change context being warmer night temperatures (Tougeron & Sanders 2023). A recent study revealed that higher temperatures increased top-down control of the predatory *Mesostoma* flatworms on *Daphnia*, changing the zooplankton community structure and affecting algal biomass (Devkota et al. 2023). The small volume of the urban habitats from which we isolated the flatworms (Tüzün et al. 2025) makes *Daphnia* particularly susceptible to the urban heat island effect. Therefore, we suggest further exploration of the interaction between warming, ALAN and flatworm parasitism on *Daphnia* population dynamics, zooplankton community composition and top-down control of phytoplankton in urban ponds.

## ACKNOWLEDGMENT

We greatly appreciate the assistance of Johannes Reichenbach during the experiment. NT was supported by the Alexander von Humboldt Research Fellowship and Marie Skłodowska-Curie Actions Fellowship.

## DATA AVAILABILITY STATEMENT

Data from the experimental trial have been deposited at FigShare: https://doi.org/10.6084/m9.figshare.28457171

## SUPPLEMENTARY MATERIAL

**Table S1.** Parameter estimates (with 95% confidence intervals) of logistic growth models, shown for the two light pollution treatment groups of *Daphnia* that were not exposed to flatworms. Note that logistic growth models did not converge for the flatworm-treatment groups, hence no estimates are provided. Overlapping confidence intervals around the estimates indicate no significant difference between light pollution treatments.

| Light pollution treatment | Carrying capacity | Time at inflection point | Exponential growth |
| --- | --- | --- | --- |
| Dark at night | 50.63 (42.96 - 67.55) | 11.08 (9.59 - 13.34) | 1.88 (0.77 - 4.28) |
| Light at night | 58.91 (49.24 - 93.80) | 12.14 (10.89 - 16.39) | 2.43 (1.08 - 4.85) |

**Figure S1.**
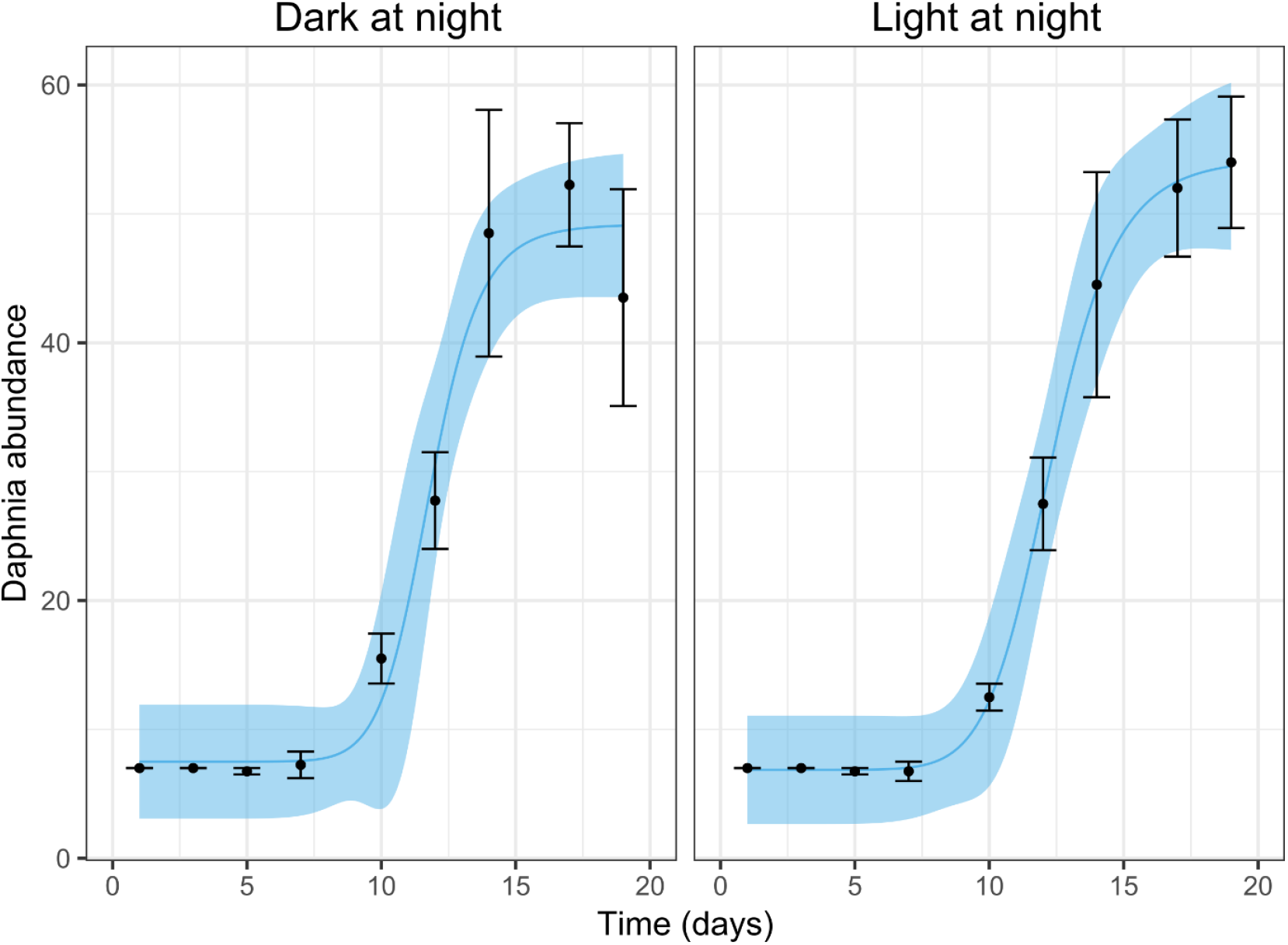
*Daphnia* abundance over time as a function of the presence or absence of light pollution at night. Left panel represents no light pollution (i.e. dark at night) condition, right panel represents light pollution condition. Shown are logistic growth curves for the control-treatment (note that flatworm-treatment was ignored here, as the data do not follow a logistic growth function; see Fig.2). Shown are also raw means (± 1SE) per time point. Bands around lines represent 95% confidence intervals (derived from the logistic growth model).

